# Retroviruses integrate into a shared, non-palindromic motif

**DOI:** 10.1101/034991

**Authors:** Paul D. W. Kirk, Maxime Huvet, Anat Melamed, Goedele N. Maertens, Charles R. M. Bangham

**Affiliations:** MRC Biostatistics Unit, Cambridge Institute for Public Health, Cambridge, UK; Centre for Integrative Systems Biology and Bioinformatics, Department of Life Sciences, Imperial College London, London, UK; Section of Virology, Division of Infectious Diseases, Imperial College London, London, UK

## Abstract

Palindromic consensus nucleotide sequences are found at the genomic integration sites of retroviruses and other transposable elements. It has been suggested that the palindromic consensus arises as a consequence of structural symmetry in the integrase complex, but the precise mechanism has yet to be elucidated. Here we perform a statistical analysis of large datasets of HTLV-1 and HIV-1 integration sites. The results show that the palindromic consensus sequence is not present in individual integration sites, but appears to arise in the population average as a consequence of the existence of a non-palindromic nucleotide motif that occurs in approximately equal proportions on the plus-strand and the minus-strand of the host genome. We demonstrate that palindromic probability position matrices are characteristic of such situations. We develop a generally applicable algorithm to sort the individual integration site sequences into plus-strand and minus-strand subpopulations. We apply this algorithm to identify the respective integration site nucleotide motifs of five retroviruses of different genera: HTLV-1, HIV-1, MLV, ASLV, and PFV. The results reveal a non-palindromic motif that is shared between these retroviruses.

## 1 Introduction

Integration of a copy of the viral RNA genome is essential to establish infection by retroviruses. This process (see, for example,^1, 2^ for reviews) is catalyzed by the virally encoded enzyme integrase (IN) and is composed of two steps: (i) the 3’ processing reaction; and (ii) strand transfer. During the 3’ processing reaction, a di- or tri-nucleotide is removed from the 3’ ends of the viral long terminal repeats (LTRs) to expose the nucleophilic 3’OH groups that consequently attack the phosphodiester backbone of the target DNA during strand transfer. Strand transfer results in single stranded DNA gaps that are filled in and repaired by host cellular enzymes. Depending on the retrovirus, the strand transfer reaction takes place with a 4 (e.g. MLV and prototype foamy virus, PFV), 5 (e.g. HIV-1) or 6 (e.g. HTLV-1 and 2) base pair stagger, giving rise to equally numbered target duplication sites.

Integration is not random: each retrovirus has characteristic preferences for the genomic integration site (IS) (e.g.^3–9^). These preferences are evident on at least three scales: chromatin conformation and intranuclear location; proximity to specific genomic features such as transcription start sites or transcription factor binding sites; and the primary DNA sequence at the IS itself. Certain host factors also play an active part in determining the genomic integration site. The best characterized of such factors are LEDGF,^10, 11^ which biases HIV-1 integration into genes in preference to intergenic regions,^12^ and BET proteins, which direct MLV integration into the 5’ end of genes.^13–15^

At the DNA sequence level, previous studies have revealed a weak palindromic consensus sequence at the IS in several retroviral infections, including HTLV-1, ASLV, FV, MLV, SIV, and HIV-1.^16–18^ The reason for the presence of a palindromic consensus sequence remains unknown, but authors have speculated that it reflects the binding to the DNA of the pre-integration complex (PIC) in symmetrical dimers or tetramers, so that each half-complex has a similar DNA target preference.^16^ However, the consensus sequence is a population average, defined by taking the modal nucleotide at each position in a population of IS sequences. The question arises whether or not the consensus is truly representative of the population. It may be a poor representation of the population if, for example, the population exhibits a high degree of variability, or if the population is composed of two or more distinct subpopulations (and hence is bi- or multi-modal). It is known that retroviral IS sequences are highly diverse, which immediately indicates the need for caution when interpreting the consensus. Here we perform statistical analyses to determine whether or not the palindromic consensus sequences are efficient representative summaries of the populations of IS sequences from which they are calculated. We find strong evidence that this is not the case, and investigate the possibility that these palindromic consensus sequences arise from the presence of motif sequences that appear in both “forward” and “reverse complement” orientations in the genome.

### 1.1 Palindromic sequences and PPMs

In everyday usage, a palindrome is a word that reads the same forwards as backwards. Palindromes may have an even number of letters (e.g. HANNAH), or an odd number (e.g. KAYAK). The axis of symmetry lies between the letters in an even palindrome (HAN | NAH), but passes through the central letter in an odd palindrome (KAYAK). The definition of a palindrome for a nucleotide sequence is slightly different: a nucleotide sequence is said to be palindromic if it is equal to its reverse complement (e.g. GAATTC and its complement, CTTAAG). Odd palindromes are possible in nucleotide sequences provided we allow “or” operators; e.g. GAA[CG]TTC, where [CG] indicates “either a C or a G”.

When dealing with multiple, aligned sequences, it is useful to define the notion of a palindromic *position probability matrix* (PPM). Given a collection of *N* aligned DNA sequences each of length *q*, the PPM for the collection is a matrix *P*, of size 4 × *q* whose rows are indexed by the letters A, T, C, G, and whose columns are indexed by the positions 1,…,*q*. The (*i, j*)-entry of the PPM is the estimated marginal probability of observing letter *i* in position *j* of any sequence in our collection. We may also define the *reverse complement* PPM *P*^(^*^RC^*^)^, which is the PPM for the collection of reverse complement sequences, and may be obtained from *P* by first swapping the ‘A’ and ‘T’ rows, and also the ‘C’ and ‘G’ rows, and then reversing the order of the columns (see Supplementary Methods). We define a PPM to be palindromic if *P = P*^(^*^RC^*^)^. A palindromic PPM may be either even or odd, according to the parity of *q*. Note that a collection of sequences that has a palindromic PPM necessarily has a palindromic consensus sequence, but the converse does not hold (i.e. if a collection of sequences possesses a palindromic consensus, they may or may not have a palindromic PPM). The target integration sites for HTLV-1 and HIV-1 not only possess palindromic consensus sequences, but also palindromic PPMs (Fig. 1).

**Figure 1.**
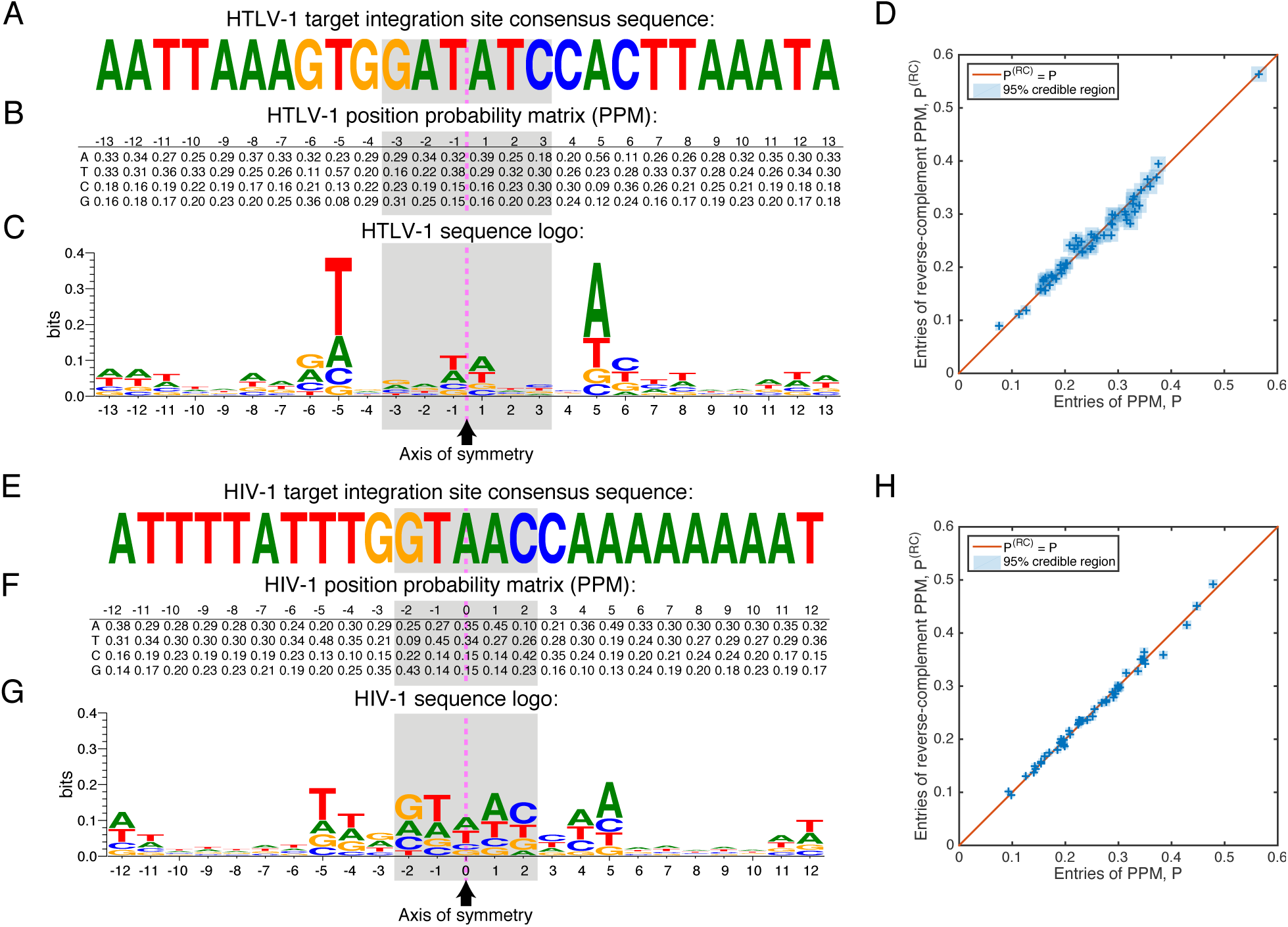
Palindromic HTLV-1 and HIV-1 target integration site consensus sequences and PPMs, calculated from 4,521 HTLV-1 and 13,442 HIV-1 IS sequences. (A) In agreement with previous studies, we find the HTLV-1 consensus sequence to be a distinctive weak palindrome. The palindrome’s axis of symmetry is indicated by the dashed pink line, while the shaded region indicates the duplicated region. (B) The PPM, *P*, for the target integration sites is also palindromic, i.e. *P*_1_*_,−j_* ≈ *P*_2_*_,j_*, *P*_2_*_,−j_* ≈ *P*_1_*_,j,_ P*_3_*_,−j_* ≈ *P*_4_*_,j_*, and *P*_4_*_,−j_* ≈ *P*_3_*_,j_* for *j* = 1,…, 13. Sequence positions to the left of the symmetry line are labeled as negative, and those to the right as positive. (C) The symmetry in the PPM may be conveniently visualized using a sequence logo, which also highlights that the palindrome is only weak (has low information content). (D) We plot the entries in the first 13 columns of the PPM, *P*, against the corresponding entries in the reverse-complement PPM, *P*^(^*^RC^*^)^, i.e. 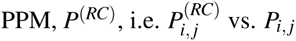. Uncertainty in the PPM entries is indicated using blue squares showing the 95% credible interval (highest posterior density) range (see Supplementary Methods). A perfect palindromic PPM would be one for which *P*^(^*^RC^*^)^ *= P*, whose entries would lie along the diagonal shown in the plot. (E) – (F): As (A) – (D), but using the HIV-1 integration sites.

### 1.2 How palindromic PPMs may arise

The presence of palindromic PPMs at the target integration sites of HTLV-1 and HIV-1 is anomalous. While symmetry in the integrase complex might be a plausible explanation for the existence of palindromic consensus sequences, it is not clear how such symmetry could generate a palindromic PPM, since a PPM provides a summary description of the proportions of *all* 4 nucleotides at *all q* sequence positions — not just the most frequently appearing nucleotides, and not just at a few positions. If the symmetry in the integrase complex is not a likely explanation for the palindromic PPMs, how else might they have arisen?

Suppose we have a collection of 2*N* sequences of length *q*, with corresponding PPM denoted by *P*, which is non-palindromic. If we randomly replace *N* of these sequences with their reverse complement sequences, the PPM for the resulting collection will be 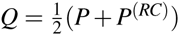. It is straightforward to show that 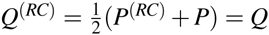, and hence *Q* is palindromic. Thus, given any collection of sequences, we can always construct a new collection that possesses a palindromic PPM (and hence a palindromic consensus) by randomly taking the reverse complement of half the sequences. Given a collection of sequences that has a palindromic PPM, *Q* (such as the retroviral target integration sites), it therefore seems sensible to ask if *Q* is truly representative of a typical individual sequence of the collection (which we will refer to as the *true palindrome* hypothesis); or if instead our sequences fall into two approximately equally sized populations, one characterized by a (non-palindromic) PPM P, and the other characterized by the reverse complement PPM, *P*^(^*^RC^*^)^. We will refer to the second possibility, i.e. the hypothesis that the consensus arises from a mixture of two complementary motifs, as the *complementary subpopulations* hypothesis. If this hypothesis is true, we would need to “un-mix” our collection of sequences into its two constituent subpopulations in order to identify the PPM, *P*, that truly characterizes the collection, i.e. the motif that characterizes the preferred integrase target site.

### 1.3 No evidence of palindrome within individual sequences

Let **s** = σ_1_σ_2_…σ_*q*−1_σ_*q*_ denote a sequence of length *q*, where σ_*i*_ ∈ {*A, T, C, G*} for all *i*. When dealing with palindromes, it is convenient to relabel the indices to reflect the location of the axis of symmetry. We will assume, without loss of generality, that the axis of symmetry is at the center of the consensus sequence, in which case we may relabel the indices so that: s = σ_−_*_n_*σ_−_*_n+_*_1_…σ_−2_σ*_−_*_1_σ*_+_*_1_ *…* σ*_+n−_*_1_σ*_+n_* if *q* = 2*n* is even; or s = σ_−_*_n_*σ_−_*_n+_*_1_… σ_−2_σ*_−_*_1_σ*_+_*_1_σ_0_σ*_+_*_1_ *…* σ*_+n−_*_1_σ*_+n_* if *q = 2n +* 1 is odd. Using this notation, it follows that if s is a perfect palindrome, then (σ*_−i_*, σ*_+i_*) is a complementary pair — e.g. (A,T) or (C,G) — for all *i =* 1,…,*n*.

Let s^(1)^,…, s^(^*^N^*^)^ be the collection of IS sequences used to calculate palindromic consensus sequence s^(CON)^. We wish to assess how “close” each sequence is to being palindromic. We therefore define the *palindrome index* (PI) for sequence s, denoted *ρ* (s), to be the proportion of positions at which (σ*_−i_*, σ*_+i_*) is a complementary pair (see Materials and Methods). It follows that the palindrome index is 1 if s is perfectly palindromic, and 0 if there are no positions at which s matches its reverse complement.

Note that, by chance, we would expect *ρ* (s) to be higher for shorter sequences (since, for example, the probability of a random sequence of length 2 being perfectly palindromic is clearly much greater than the probability for a random sequence of length 16). For this reason, and as is common for indices such as this (e.g. the Rand Index^19,20^), we also introduce a “corrected for chance” version of the palindrome index (see Materials and Methods). We denote this by *ρ_A_* and refer to it as the *adjusted palindrome index* (API). The API is 1 if the sequence is perfectly palindromic, 0 if the sequence is as palindromic as expected by chance, and negative if the sequence is *less* palindromic than expected by chance.

We calculated the PI and API for each of the HTLV-1 and HIV-1 IS sequences, as well as for the consensus sequences. We considered a range of sequence lengths: 2*n* = 2,4,…, 26 for HTLV-1, and (2*n* +1) = 3,5,…, 25 for HIV-1. Results are shown in Table 1 (for HTLV-1) and Table 2 (for HIV-1), and illustrated in Figure 2.

**Table 1.** Palindrome index (PI) and adjusted palindrome index (API) scores for HTLV-1 integration site sequences. The mean PI values were calculated by finding the PI for each of the 4,521 individual IS sequences, and then taking the mean (similarly for the mean API values).

| Seq.<br>length,<br>$2n$ | PI for<br>consensus,<br>$\rho(s^{(\text{CON})})$ | Mean<br>$PI$,<br>$\bar{\rho}$ | API for<br>consensus,<br>$\rho_A(s^{(\text{CON})})$ | Mean<br>API,<br>$\bar{\rho}_A$ | $p$-value<br>($\mathcal{H}_0 : \bar{\rho}_A = 0$) |
| --- | --- | --- | --- | --- | --- |
| 26 | 0.85 | 0.27 | 0.79 | -0.01 | 2.12E-06 |
| 24 | 0.92 | 0.27 | 0.89 | -0.01 | 2.99E-07 |
| 22 | 0.91 | 0.27 | 0.87 | -0.01 | 5.31E-07 |
| 20 | 0.90 | 0.27 | 0.86 | -0.02 | 1.58E-07 |
| 18 | 0.89 | 0.27 | 0.85 | -0.02 | 1.08E-07 |
| 16 | 1.00 | 0.27 | 1.00 | -0.02 | 2.41E-11 |
| 14 | 1.00 | 0.27 | 1.00 | -0.03 | 5.00E-15 |
| 12 | 1.00 | 0.27 | 1.00 | -0.03 | 1.08E-14 |
| 10 | 1.00 | 0.27 | 1.00 | -0.04 | 1.58E-18 |
| 8 | 1.00 | 0.24 | 1.00 | -0.03 | 1.15E-14 |
| 6 | 1.00 | 0.24 | 1.00 | -0.04 | 5.04E-18 |
| 4 | 1.00 | 0.24 | 1.00 | -0.05 | 1.28E-15 |
| 2 | 1.00 | 0.23 | 1.00 | -0.08 | 2.83E-21 |

**Table 2.** Palindrome index (PI) and adjusted palindrome index (API) scores for HIV-1 integration site sequences.

| Seq.<br>length,<br>$2n + 1$ | PI for<br>consensus,<br>$\rho(s^{(\text{CON})})$ | Mean<br>$PI$,<br>$\bar{\rho}$ | API for<br>consensus,<br>$\rho_A(s^{(\text{CON})})$ | Mean<br>API,<br>$\bar{\rho}_A$ | $p$-value<br>($\mathcal{H}_0 : \bar{\rho}_A = 0$) |
| --- | --- | --- | --- | --- | --- |
| 25 | 0.92 | 0.27 | 0.88 | -0.01 | 8.21E-09 |
| 23 | 0.91 | 0.27 | 0.87 | -0.01 | 1.60E-08 |
| 21 | 0.90 | 0.27 | 0.86 | -0.01 | 4.29E-09 |
| 19 | 0.89 | 0.27 | 0.85 | -0.01 | 1.29E-11 |
| 17 | 0.88 | 0.27 | 0.83 | -0.01 | 1.08E-12 |
| 15 | 0.86 | 0.28 | 0.80 | -0.02 | 1.04E-13 |
| 13 | 1.00 | 0.28 | 1.00 | -0.02 | 3.16E-18 |
| 11 | 1.00 | 0.28 | 1.00 | -0.03 | 1.69E-26 |
| 9 | 1.00 | 0.27 | 1.00 | -0.03 | 1.02E-27 |
| 7 | 1.00 | 0.27 | 1.00 | -0.03 | 8.57E-25 |
| 5 | 1.00 | 0.28 | 1.00 | -0.04 | 1.09E-24 |
| 3 | 1.00 | 0.27 | 1.00 | -0.07 | 1.95E-35 |

**Figure 2.**
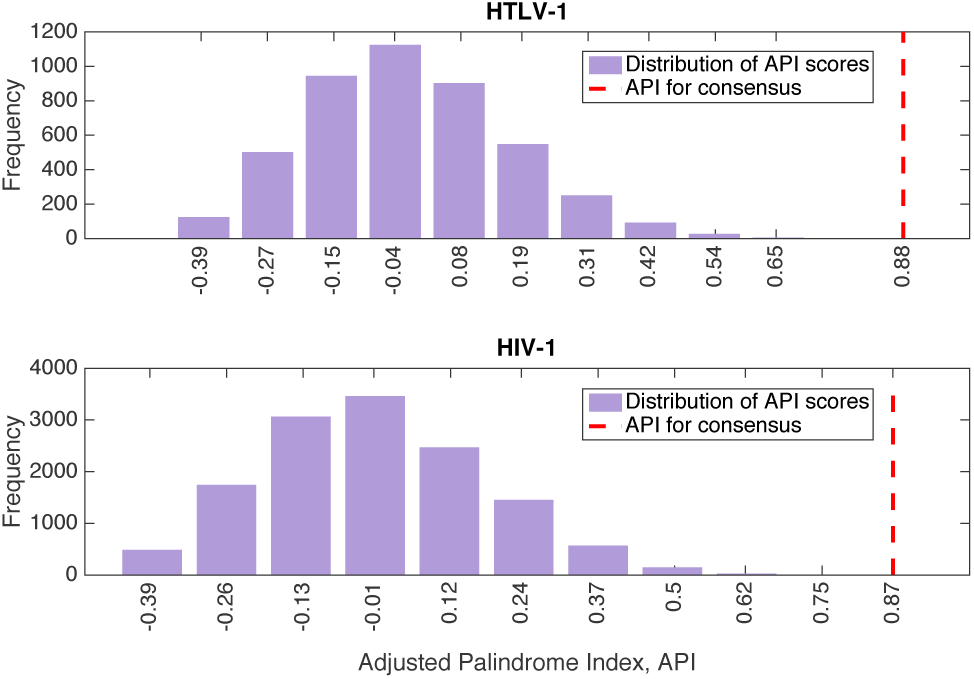
Distribution of adjusted palindrome index (API) scores over all 4,521 HTLV-1 integration site sequences (top, taking the sequence length to be 2*n* = 26), and over all 13,442 HIV-1 integration sequences (bottom, with 2*n* +1 = 25). In both cases, the API for the corresponding consensus sequence (indicated by the red dashed line) is in the extreme positive tail of the distribution.

For both the HTLV-1 and HIV-1 sequences, and for all values of n, the API (and PI) scores for the consensus sequences are much higher than the mean API (respectively PI) scores calculated over all the individual IS sequences. This observation is consistent with Figure 2: for both the HTLV-1 and HIV-1 IS sequences, the API for the consensus sequence is in the extreme right tail of the distribution of API scores. The consensus sequences are therefore much more palindromic than the individual IS sequences from which they were derived.

Moreover, the mean API scores for both the HTLV-1 and HIV-1 IS sequences are consistently negative, for all values of *n*. Although small in magnitude, one-sample *t*-tests confirmed that these values were significantly different from zero (*p*-values provided in the final columns of Tables 1 and 2). This indicates that the individual IS sequences are, on average, slightly *less* palindromic than we would expect by chance alone. This result is consistent with the existence of a non-palindromic motif. We conclude that there is no evidence of a palindrome within individual IS sequences.

### 1.4 Probability model for the IS sequences

We next test the *complementary subpopulations* hypothesis. This hypothesis makes the following predictions: (i) the collection of sequences can be split into two subcollections described by reverse complementary PPMs, *P* and *P*^(^*^RC^*^)^ with *P ≠ P*^(^*^RC^*^)^; and (ii) this split provides a better description of the data than that provided by a single palindromic PPM *Q*.

The natural way in which to identify subpopulations is to model the collection of sequences using a 2-component mixture model, with one component corresponding to the subpopulation of sequences in the “forward” orientation and the other corresponding to those in the “reverse complement” orientation. Note that the labels “forward” and “reverse complement” are strictly relative descriptions. Our model is thus,

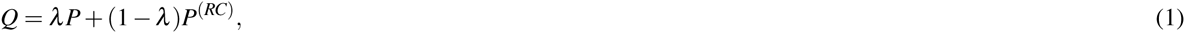

where *Q* is the overall (palindromic) PPM for the whole population of sequences, and *λ* ∈ [0, 1] is the proportion of sequences belonging to the subcollection with PPM *P*. Fixing *λ* = 0.5 would be consistent with the complementary subpopulations hypothesis; however, we treat *λ* as a free parameter, i.e. to be estimated from the data rather than introducing an assumption. Note further that the model includes the possibility of a true palindrome (i.e. that the data are best described as a single population with a palindromic PPM) as a special case, when *P = P*^(^*^RC)^*.

We implemented several algorithms for fitting the mixture model (see Supplementary Methods). Here we present the results obtained using the expectation-maximization (EM) algorithm^21^ to find maximum likelihood estimates for the model’s parameters (algorithm details provided in Materials and Methods).

## 2 MATERIALS AND METHODS

### 2.1 Mapped integration sites

To focus on the initial integration targeting profile of HTLV-1 and HIV-1, integration sites were identified in DNA purified from cells infected experimentally *in vitro*. Jurkat T-cells were infected either by short co-culture with HTLV-1-producing cell line MT2^22^ or by VSV-G pseudotyped HIV-1 (kind gift from Dr. Ariberto Fassati, UCL). Identification of 4,521 HTLV-1 integration sites from *in vitro* infected Jurkat T-cells has been described before.^7,23^ Identification of 13,442 HIV-1 integration sites was carried out using a similar approach, using the following HIV-specific PCR forward primers: HIVB3 5’-GCTTGCCTTGAGTGCTTCAAGTAGTGTG-3’, HIVP5B5 5’-AATGATACGGCGACCACCGAGATCTACACGTGCCCGTCTGTTGTGTGACTCTGG-3’ and HIV-specific sequencing primer 5’-ATCCCTCAGACCCTTTTAGTCAGTGTGGAAAA TCTC-3’.

### 2.2 Palindrome index (PI)

We denote by s^(CON)^ the consensus sequence calculated from sequences s^(1)^,…,s^(^*^N^*^)^, and suppose that s^(CON)^ is approximately palindromic, with axis of symmetry at the center of the sequence. If the palindrome is initially odd, we remove the central letter from all sequences, so that the palindrome becomes even. It follows that all sequences may be assumed to be of even length, 2*n*, and hence may be written 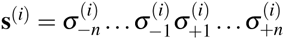 for *i =* 1,…,*N*. We define the palindrome index, *ρ*, for the sequence s^(^*^i^*^)^ to be 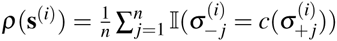, where *c*(*X*) denotes the complement of *X* (e.g. *c*(*A*) *= T*) and 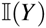 denotes the indicator function, which is 1 if *Y* is true and 0 otherwise. The palindrome index is therefore the proportion of positions at which s^(^*^i^*^)^ is equal to its reverse complement.

### 2.3 Adjusted palindrome index (API)

By chance alone, the palindrome index will tend to be larger for smaller values of *n* than for larger values. We therefore introduce a “corrected for chance” version of the index (see, for example,^24^) to account for this: 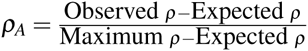. The maximum value for *ρ* is 1 (when a sequence is perfectly palindromic). The expected value for *ρ* is the expectation when 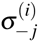 and 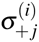 are independent, 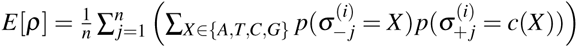, where the probabilities 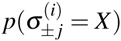 are the empirical marginal probabilities, which may be taken from the entries of the PPM (e.g. Figure 1B and 1F).

### 2.4 Two component mixture model

We model the IS sequences as being drawn from a 2-component mixture model, *p*(s|*P*,λ) = λ*f* (s|*P*) + (1 − *λ*)*f* (s|*P*^(^*^RC^*^)^), where *f*(s|*P*) is the likelihood of sequence s given PPM *P*. The likelihood is straightforwardly defined as the product of probabilities of each of the elements of s, 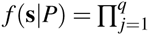 Prob(letter *σ_j_* in position *j* | *P*), where the individual probabilities are given by the entries of the PPM.

### 2.5 Expectation-maximization (EM) algorithm for our model

We refer the reader to^21^’^25^ for general information about the EM algorithm, and here provide the update equations for the model parameters, *λ* and *P*. At iteration *t*, define 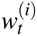 to be the posterior probability of sequence s^(^*^i^*^)^ belonging to the subpopulation with PPM *P*, given *λ_t_*_−1_ and *P_t_*_−1_ (the parameter estimates at iteration *t −* 1). That is,

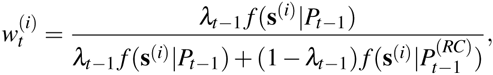

where *f* (s|*P*) is as defined previously. Also, for *X* ∈ {*A, T, C, G*} and *k* = 1,…, *n* (or *k* = 0,…, *n*, in the odd palindrome case), define

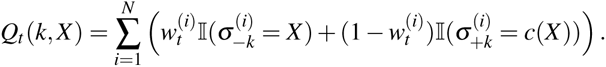

Then 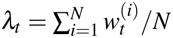, and defining the element of *P_t_* in column *k* and row labeled by nucleotide *X* to be *P_t_* (*k, X*), we have:

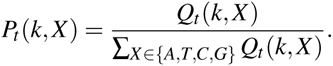

Initialization details and stopping criteria are provided in Supplementary Methods.

## 3 RESULTS

All considered algorithms for fitting the mixture models provided qualitatively identical results (see Supplementary Results). For both HTLV-1 and HIV-1, the algorithms identified complementary subpopulations within the collections of IS sequences (Fig. 3A), with the subpopulations appearing in approximately equal proportions (*λ_HTLV_* = 0.47 and *λ_HIV_* = 0.49).

**Figure 3.**
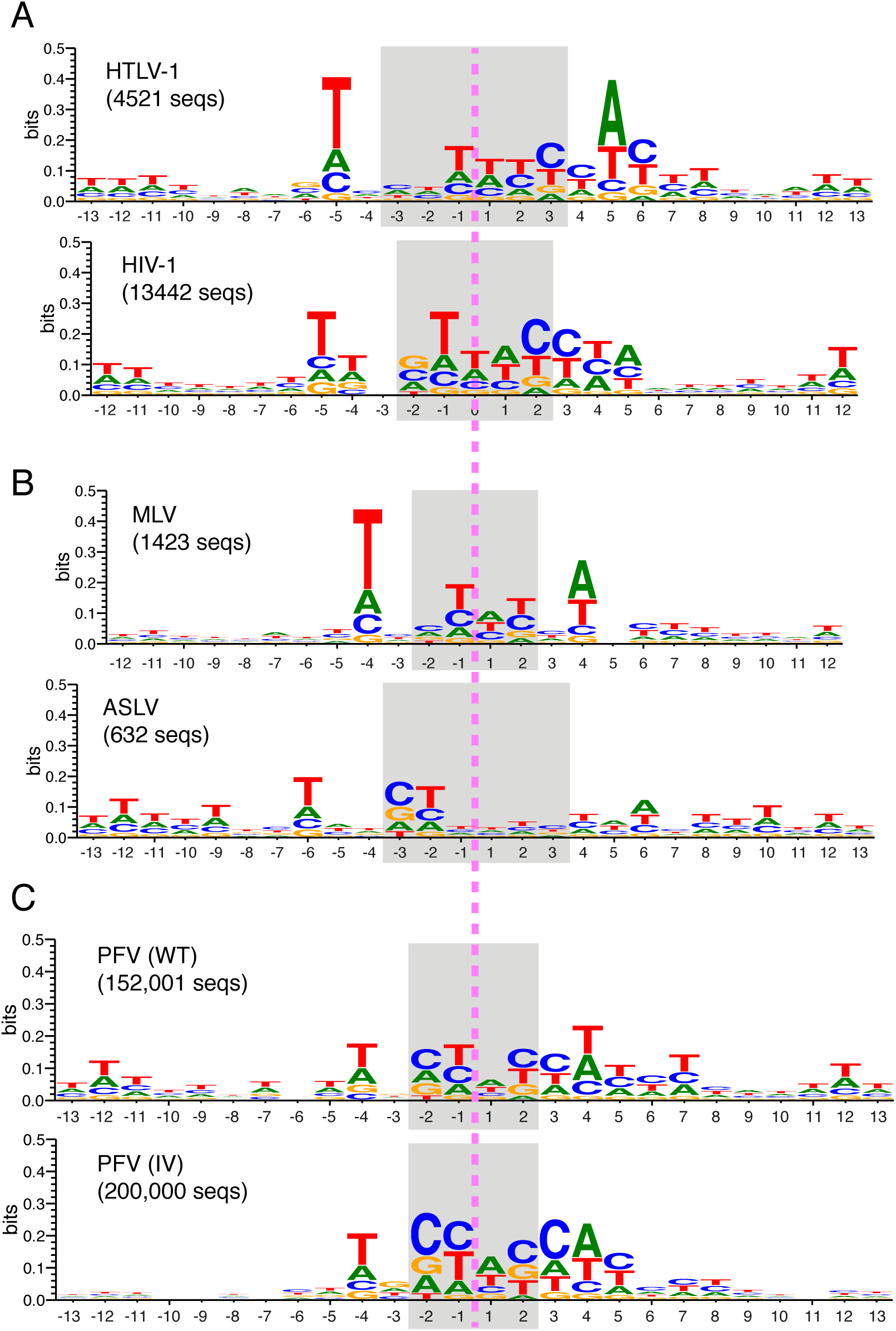
Summary of results from applying the EM algorithm to fit the 2-component mixture model. (A) Sequence logo summaries of one of the two subpopulations of integration site sequences in the HTLV-1 and HIV-1 datasets (in each case, the other subpopulation is characterized by the reverse complement of the sequence logo shown). (B) As (A), but for the MLV and ASLV datasets. (C) As (A), but for the PFV (WT) and PFV (IV) datasets.

### 3.1 Quality of fit

We next assessed whether the complementary subpopulations hypothesis provided a better description of the data than the true palindrome hypothesis. Note that, since *P*^(^*^RC^*^)^ is completely determined by *P*, the 2-component mixture model has only one more free parameter (λ) than the case where we assume the data are best described as a single population.

Although it is tempting to apply a simple likelihood ratio test (LRT) to determine if the unconstrained model provides a significantly better fit to the data than the constrained, true palindrome model (in which *P* = *P*^(^*^RC^*^)^), it is well known that for mixture models the LRT statistic does not in general follow standard χ^2^ distributions.^26^ We therefore adopted McLachlan’s approach^27^ in order to construct an empirical null distribution for the LRT statistic, *D*. Note that here the null model is a single component with PPM equal to the empirical PPM (given in Figure 1B for HTLV-1 and Figure 1F for HIV-1), while the alternative is the fitted 2-component mixture model. Briefly, we simulated 1,000 new datasets using the null model, fitted both the null and alternative models to each simulated dataset, and calculated the LRT statistic each time. In this way, we obtained empirical null distributions for the LRT statistic, which we then used to assess the significance of the observed LRT statistics. For the HTLV-1 IS sequences, the 1,000 values sampled from the null distribution of the LRT statistic all fell between −28.64 and 18.79, while the observed LRT statistic was 1.49 *×* 10^3^. For the HIV-1 IS sequences, the sampled LRT statistics all fell between −32.37 and 29.24, while the observed LRT statistic was 2.86 *×* 10^3^. For both the HTLV-1 and HIV-1 datasets we may clearly reject the null model in favour of the alternative model (*p* < 0.001).

For both datasets, we also calculated the difference in Bayesian Information Criterion (ΔBIC) statistic for the two competing models, giving ΔBIC_HIV_ = 2.86 *×* 10^3^ and ΔBIC_HTLV_ = 1.48 *×* 10^3^. Assuming ΔBIC provides an adequate approximation to twice the natural logarithm of the Bayes Factor,^28^ any value of ΔBIC greater than 10 would be interpreted as providing *very strong* evidence against the null, one-population model.^29^

### 3.2 Other retroviruses

We additionally fitted our 2-component mixture model to smaller datasets on HTLV-1, HIV-1, MLV, and ASLV taken from the literature.^17^ While the reduced sample sizes limit confidence in the inferences drawn from these datasets, they may nevertheless provide useful qualitative results. The results on MLV and ASLV are given in Fig. 3B: the results on HTLV-1 and HIV-1 are qualitatively identical to those obtained from the larger datasets, and are given in Supplementary Results. We also considered two large PFV datasets from:^30^ (i) the PFV (WT) dataset, which comprises integration sites for 153,447 unique integration events in HT1080 cells; and (ii) the PFV (IV) dataset, comprising approximately 2 *×* 10^6^ integration sites determined using purified PFV intasomes and deproteinised human DNA. After pre-processing to remove duplicates and sequences containing Ns, 152,001 integration sites remained in the PFV (WT) dataset and 2,197,613 in the PFV (IV) dataset. To reduce computation time, we randomly sampled 200,000 integration site sequences from the PFV (IV) dataset to use for analysis. The results on PFV (WT) and PFV (IV) are given in Fig. 3C. The results obtained for all retroviruses have a number of remarkable similarities (see Discussion).

To assess the implications of our results for the flexibility properties of retroviral integration sites, we calculated dinucleotide pyrimidine-purine (YR) and purine-pyrimidine (RY) profiles^31^ for the collection of sequences in each subpopulation, which we found to be similar to those calculated for the overall population (see Supplementary Results).

## 4 DISCUSSION

The factors that influence the pattern of integration of retroviruses and transposable elements operate at different physical scales. The strength of association between specific genomic features and retroviral integration frequency depends on the genomic scale on which the data are analysed.^32, 33^ Broadly, three scales have been studied: chromosome domains and euchromatin/heterochromatin; genomic features such as histone modifications and transcription factor binding sites; and primary DNA sequence. At the largest scale (chromosome domains and heterochromatin/euchromatin), it was recently demonstrated that HIV-1 integration is biased towards regions of chromatin that lie in proximity to the nuclear pores, through which the HIV-1 preintegration complex enters the nucleus.^8,9^

The primary DNA sequence of the host genome is thought to influence the site of retroviral integration by determining both the binding affinity of the intasome and the physical characteristics of the target DNA, especially the ability of the double helix to bend,^34,35^ which depends in turn on the presence of specific dinucleotides and trinucleotides. Muller and Varmus^36^ concluded that the bendability of DNA could explain the preferential integration of certain retroviruses in DNA associated with nucleosomes. The requirement for DNA bending during retroviral integration has been explained by the discovery of the crystal structure of the foamy viral intasome complexed with target DNA.^31,37^ Complete unstacking of the central dinucleotide at the site of integration allows the scissile phosphodiester backbone to reach the active sites of the IN protomers.^37^ Although the bending of the tDNA observed in the crystal structure does not correspond with the bend described in nucleosomal DNA,^38^ the EM structure of the foamy viral intasome in complex with mononucleosomes^30^ showed that the nucleosomal DNA is lifted from the histone octamer to allow proper accommodation within the active sites of the IN protomers. Given that integration catalyzed by different retroviral INs gives rise to a different target duplication size, it is expected that DNA bending at the site of integration will be more severe for integrations with a 4 bp target duplication compared to those with a 6 bp target duplication.^31^

Whereas some retroviruses preferentially integrate into regions of dense nucleosome packing (e.g. PFV, MLV),^30^ others prefer regions of sparse nucleosome packing (e.g. HIV, ASV;^39^). However, even in cases where nucleosome sparseness is preferred, a nucleosome at the integration site itself contributes to efficient integration.

In addition to the impact of specific dinucleotides and trinucleotides on DNA bendability, the other chief impact of primary DNA sequence on retroviral integration is the presence of a primary DNA motif, i.e. preferred nucleotides at specific positions in relation to the integration site. Palindromic DNA sequences have been reported at the insertion site of transposable elements in Drosophila,^35^ yeast^40,41^ and retroviruses.^16–18,42–44^ The presence of the palindrome has been attributed by several workers to the symmetry of the multimeric viral preintegration complex.^16,18^ However, Liao *et al*.^35^ noted that, although the palindromic pattern that they observed at the insertion site of a P transposable element in Drosophila could be discerned when as few as fifty insertion sites were aligned and averaged, the palindrome was not evident at the level of a single insertion site.

It was previously assumed that the non-appearance of the palindromic nucleotide sequence in individual retroviral integration sites was due to the fact that the palindrome was weak, i.e. poorly conserved. However, in the present study we found evidence that the palindrome was statistically significantly disfavoured at the level of individual sites: the palindrome is evident only as an average – a consensus – of the population of integration sites. We propose that the most likely explanation is that the palindrome results from a mixture of sequences that contain a non-palindromic nucleotide motif in approximately equal proportions on the plus-strand and the minus-strand of the genome.

## 5 CONCLUSION

On the hypothesis of a non-palindromic nucleotide motif in approximately equal proportions on the plus-strand and the minus-strand of the genome, we sorted the populations of sequences of several different retroviral integration sites into those with a conserved motif respectively on the plus-strand and the minus-strand of the genome. The resulting alignment revealed the putative true nucleotide motif that is recognized by the intasome in each case. Comparison of these motifs between the respective viruses showed certain similarities between the sequences (Figure 3), with a shared motif 5’-T(N1/2)[C(N0/1)T↕(W1/2)C]CW - 3’, where [ and ] represent the start and end of the duplicated region, and ↕ represents the axis of symmetry. The preference of an A (T) 2 or 3 nucleotides downstream (upstream) from the integration site was previously observed and explained by a direct contact between A and the residue at the HIV-1 IN S119 equivalent position.^31,37,45,46^ It remains to be seen whether the nucleotide composition of the remainder of the shared motif, in particular the central T-rich region, is preferred because of the flexibility of the DNA at such sequences or is due to direct contact between IN and the bases. Further structural information on lenti-, gamma-, delta- and avian retroviral intasomes is needed to answer this question.

## 6 ACKNOWLEDGEMENTS

The authors wish to thank the following individuals for providing materials: Alexander Zhyvoloup and Ariberto Fassati, Division of Infection and Immunity, University College London; and Heather Niederer, Division of Infectious Diseases, Imperial College London. Additionally, we are grateful to Laurence Game and Marian Dore, Medical Research Council Clinical Sciences Centre Genomics laboratory at Hammersmith Hospital, London, UK.

